# Fast marching to branching morphologies

**DOI:** 10.1101/2024.11.16.623917

**Authors:** Anna Kriuchechnikova, Tatiana Levdik, Alexey Brazhe

## Abstract

Many living systems self-organize into tree-like hierarchical branching patterns. How the function of these systems is shaped by their morphology, and whether it reflects optimization of any resource or cost draws a sustained interest. Applications in computational biology and biomimetics require tools to generate branching morphologies across a wide range of feature space. Here we propose a simple framework to build branching structures as converging gradient-descent paths in travel-time maps, computed with fast-marching algorithm over a stochastic speed field, with a structure-dependent feedback on local speed and path targets. We demonstrate the effects of speed field profile, feedback properties and seed density and sampling order on the resulting morphologies and their transport properties. The main utility of the framework is in its simplicity and the ability to generate realistic standalone structures in 2D in 3D, as well as tiling networks resembling astrocyte syncytium in the brain.

## Introduction

Hierarchical tree-like morphologies emerge in Nature virtually anywhere where there is a need for nutrient distribution, metabolic supply or information flow: trees, corals, blood vasculature, bronchial trees, hyphal networks to name just a few. This universality motivates an interest in the organization, efficiency and robustness of such transport networks. Computational biology and biomimetics are in need for frameworks capable to simulate various branching patterns across application domains and a a wide morphology space — from neurons to slime molds to city transportation.

Some existing models of branching morphologies are designed to connect point sources by a tree under given constraints (Chandrasekhar & Navlakha, 2019; Cuntz et al., 2010), or by growing sprouts to attraction points (Runions et al., n.d.), or focused on reproduction of stochastic tip elongation and branching under spatial cues (Hannezo et al., 2017; Ųcar et al., 2021). A different approach is taken in works building optimal flow networks from densely connected nodes, where the heavily used edges acquire more weight, while non-utilized edges die out (Corson, 2010; Durand, 2007; Ronellenfitsch & Katifori, 2019). Here we propose an algorithm to build such branching patterns based on converging gradient descent paths in the fmm-defined travel-time map.

Here we propose a framework for a practical and phenomenological construction of branching patterns rather than mechanism-based simulation of morphogenesis. The framework is based on tracing converging gradient descent paths on a dynamically changed potential surface with a feedback provided by previous paths.

Fast-marching method (Sethian, 1996) and its modifications are widely used in path planning for marine and other non-manned vehicles (Garrido et al., 2017; 2020), computation of geodesic distance between complex-shaped objects, e.g. cellular compartments in electron microscopy data (Salmon et al., 2023), and morphology reconstruction. Here we use travel-time maps, computed by fast-marching method in a feedback-updated stochastic speed field as the potential surface for gradient descent paths. The key features of the proposed framework are (i) gradient descent paths in travel-time maps, based on random speed field, (ii) connectivity hierarchy emerging from path-dependent speed field updates, and (iii) dependence of topology on path seed sampling and path target updates. We demonstrate the utility of the proposed framework in creating branching patterns with radically different topologies and demonstrate that it can be used to simulated tiling networks and 3D shapes.

## Methods

### Proposed Algorithm

The algorithm is based on following gradient descent (GD) paths in some potential field from randomly sampled seeding locations to global or local minima. The potential field is modeled as travel-time map calculated by fast-marching method, which calculates arrival times of a wavefront, originating from a levelset, for example a point in the center of the modeling space, which will serve as a target for the GD paths.

#### Fast marching method

Fast marching method solves eikonal equation to compute wavefront propagation from some wave source levelset to all other points in the given space. Scalar propagation speed can be provided as an input to the algorithm. Most simulations were done in 2D, using 512×512 matrices as modeling space and to define speed profiles. In most simulations a single point in the center of the modeling space was used as the wave source, thus all paths converged to this point. In the tiling network simulation, randomly placed points were used as wave sources, in this case the paths converged to nearest local minima of the travel time map. In the case of variable path targets, a set of pixels containing a branch with occurrence count (see below) lower than some threshold was used as a wave source. In this case the new paths converged to nearest branches with low occurrence count.

#### Stochastic speed fields

Wavefront propagation in uniform conditions would result in straight GD paths. In real systems, there are often space and trophic constraints set by the local environment, which dictate jittered branches even if the only cost would be associated with building material. Here we reproduce this a randomized speed field. In the easiest case, the speed field can be just an image with spatially uncorrelated Gaussian noise, rescaled to the interval [0…1]. A slightly more natural approach would be to use multi-scale colored noise, i.e. a weighted sum of noisy images, smoothed by convolution with a Gaussian kernel with different scale parameter values 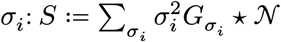, where 𝒩 is a noisy image with spatially uncorrelated normally distributed noise, ⋆ is convolution and 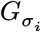 is Gaussian kernel with standard deviation σ_*i*_. Again, before applying the fast-marching method, the speed values are rescaled to [0…, 1] interval.

Some natural branching morphologies develop in an environment, where the space is locally anisotropic, e.g. due to orientations of other cells, nerve fibers, etc. We model this kind of environment as result of applying Sato’s vesselness contrast (Sato et al., 1998) to a noisy image containing spatially uncorrelated normally distributed noise. In the same way as with isotropic fields, a multiscale field can be created as a weighted sum of Sato-based contrast images with different values of the spatial scale parameter σ_*i*_.

Due to stochastic nature of the speed field, some regions can remain inaccessible to the propagating wavefront, these were excluded from building paths.

#### Converging gradient descent paths

Tree branches in the proposed algorithm are represented as GD paths, starting from some seed location and converging to the target, i.e. the fast-marching wave source. GD paths can merge, reflecting optimal routes to some locations. In the implementation, any path reaching already existing tree nodes or the path target will be stopped, and the last node of the path snippet will be linked to the tree node as a parent. The final morphology is build from these snippets. After each new path, the “path occurrence” count is updated in all affected nodes. That is, starting from the branch tip and following every parent node until the root, the counter of how many times this node participated in a path to root is updated by one. Path occurrence counts are then used in speed field updates and in path target updates if necessary.

A key aspect of the algorithm is that a new path will affect the speed profile, making it more attractive for new paths from nearby seeds to converge on the path to the tree root. In most simulations below the speed field is updated after building each path as *v*_*i*+1_ ← *v*_0_ + *O*_*i*_, where *v*_0_ is the initial speed profile and *O*_*i*_ is the occurrence count at iteration *i*. In Figure 2 we compare also logarithmic, *v*_*i*+1_ ← *v*_0_ + log(1 + *O*_*i*_), and quadratic 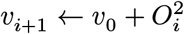 updates. Then, the travel-time map is recalculated. In the case of large trees, built from many seeds, only the first paths alter the global speed landscape, and the interval between updates of the travel-time map can be progressively increased with iteration number.

#### Seed sampling

In principle, the fast-marching wavefront will expand over whole modeling domain. For a more natural look of the presented morphologies, we limited the path seeding to the space, representing the first 50% of travel time values. For a single path target this creates a central disc with irregular boundaries, which covers around 50% of the modeling space. In the network setting, this leads to spatially overlapping domains.

Speed field updates after new paths make the algorithm greedy. This will lead to different resulting morphologies if the density or order of placing seed in simulation is different. In the results below we use either uniform seed sampling or a non-uniform sampling with central or peripheral preference. Central preference of the sampling probability is modelled as Gaussian function with σ = 75px located in the center of the 512×512 px modeling domain. Peripheral preference is modelled as the positive values of the difference between Gaussians with σ_1_ = 200px and σ_2_ = 175 px.

### Visualization of morphology patterns

The resulting morphologies are represented in two different ways, either as lines of equal width (Figure 1 – Figure 3) or with added branch diameter and brightness (Figure 4 – Figure 6). In the latter case, branch widths follow pipe flow model (Shinozaki et al., 1964), which describes how branch radius *r* of a parent node *p* is related to branch radii *r*_*i*_ at it’s children nodes 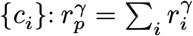; choice of values for parameter *γ* only affected morphology representation and was chosen arbitrarily. Pixel intensity at the node *n* center was scaled with node occurrence count *O*_*n*_ and decayed exponentially with distance *d* to the node center: 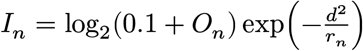, where *r*_*n*_ is branch radius at node *n*. As with the diameter, the intensity is used only for morphology representation except in Figure 6, where it was used as the speed profile for a wavefront propagation to depict signal spread in the network. The 3D structure in Figure 6 was first converted to SWC format (Parekh & Ascoli, 2013) and rendered with Shark Viewer (https://github.com/JaneliaSciComp/SharkViewer).

**Figure 1:**
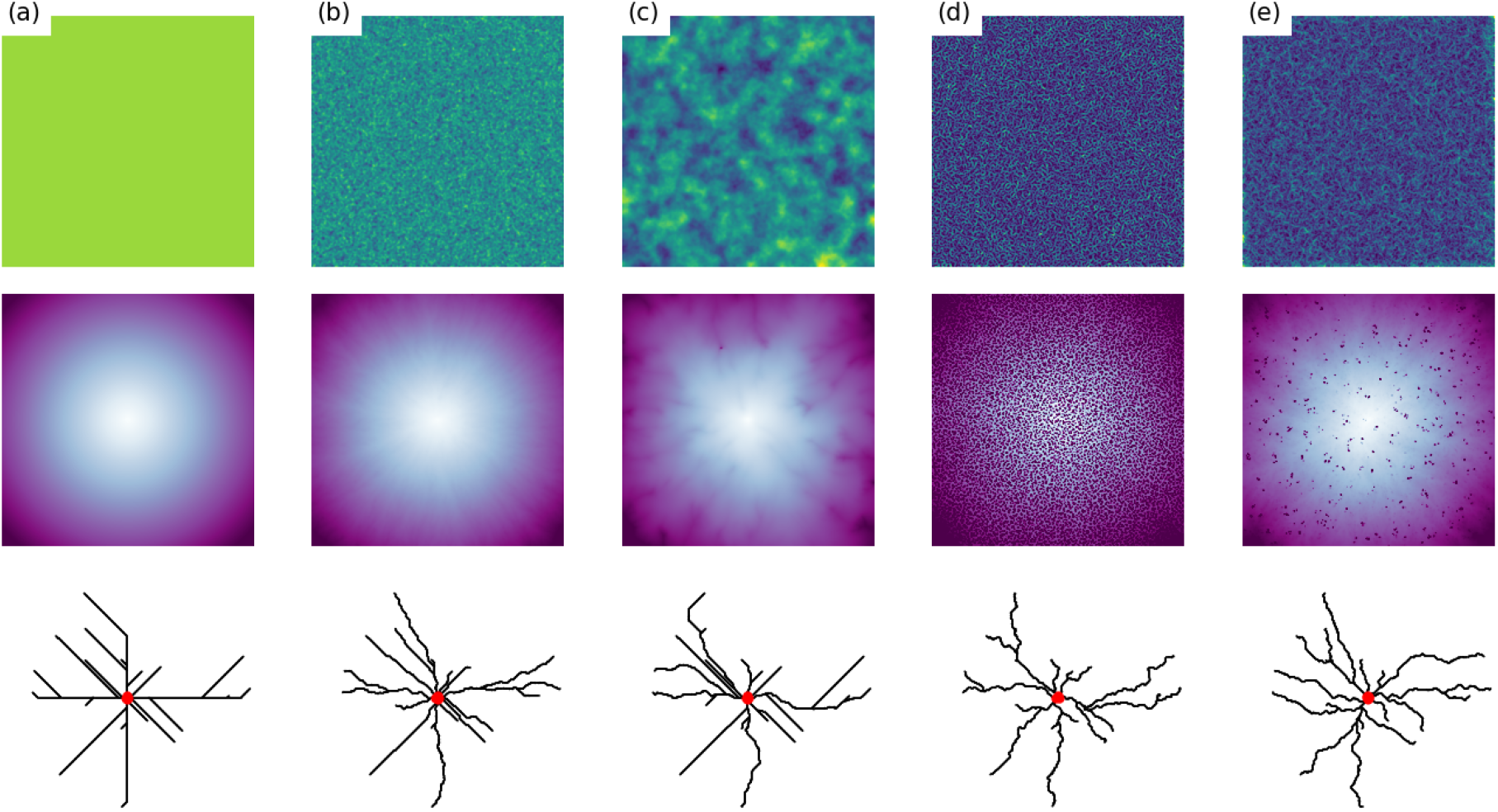
Effect of speed field on convergent gradient descent (GD) paths. Top row: speed fields, speed amplitude is show as heat map, blue—low, yellow—high; middle row: corresponding travel time maps with wave source in the center, light—short travel time, dark—long travel time; bottom row: GD paths from the same set of 20 random seed points to the source marked as red dot. In uniform field (a) travel time is smooth and the paths are straight, adding different types of noise (b-e) reshapes the travel time maps and GD paths: (b) high-pass Gaussian, (c) multi-scale Gaussian, (d) high-pass filamentous, (e) multi-scale filamentous. Note disperse points with high travel time and more jittered branches in filamentous fields.

**Figure 2:**
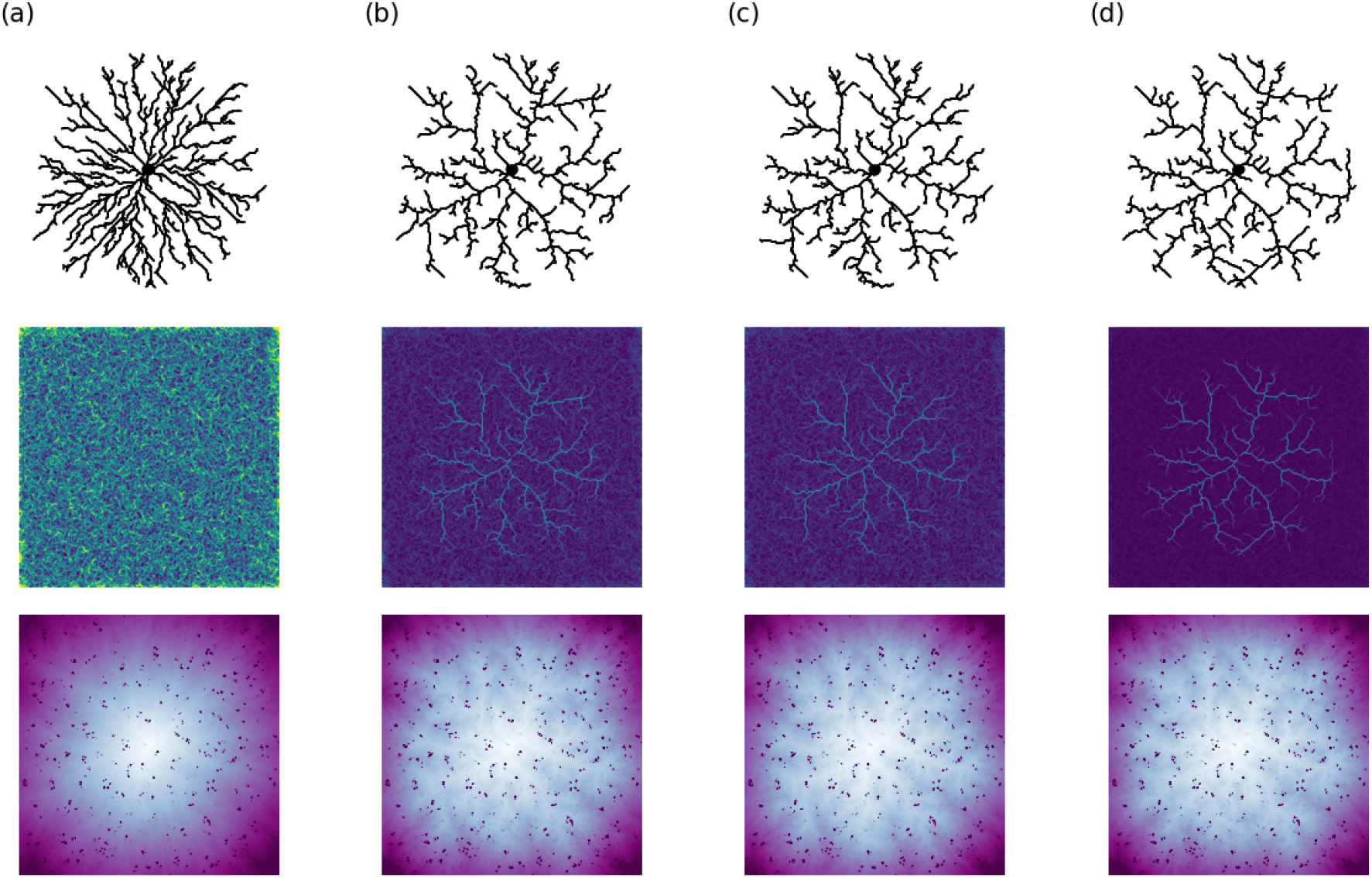
Effect of path-based feedback on velocity field, paths built from a set of randomly chosen 250 seed points and the same initial field, the one shown in (Figure 1 e). Paths follow gradient descent in travel-time maps for a wave source in the center, shown as black spot. Top row: resulting branching patterns, middle row: resulting speed field after building the tree (heat map in logarithmic intensity scale, log_2_(1 + *v*)), bottom row: resulting travel-time profiles after building the tree. (a) No action of paths on speed, *f*(*x*) → 0; speed field remains intact; (b) logarithmic scaling of path-dependent field update, *v*_*i*_ ← *v*_0_ + log_2_(*O*_*i*_), where *O*_*i*_ is the map reflecting node occurrence count in all paths (see text); (c) linear scaling *v*_*i*_ ← *v*_0_ + *O*_*i*_; (d) quadratic scaling 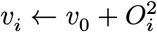. Note that with increasing nonlinearity of update, the speed profile is increasingly dominated by traces left by already built paths.

**Figure 3:**
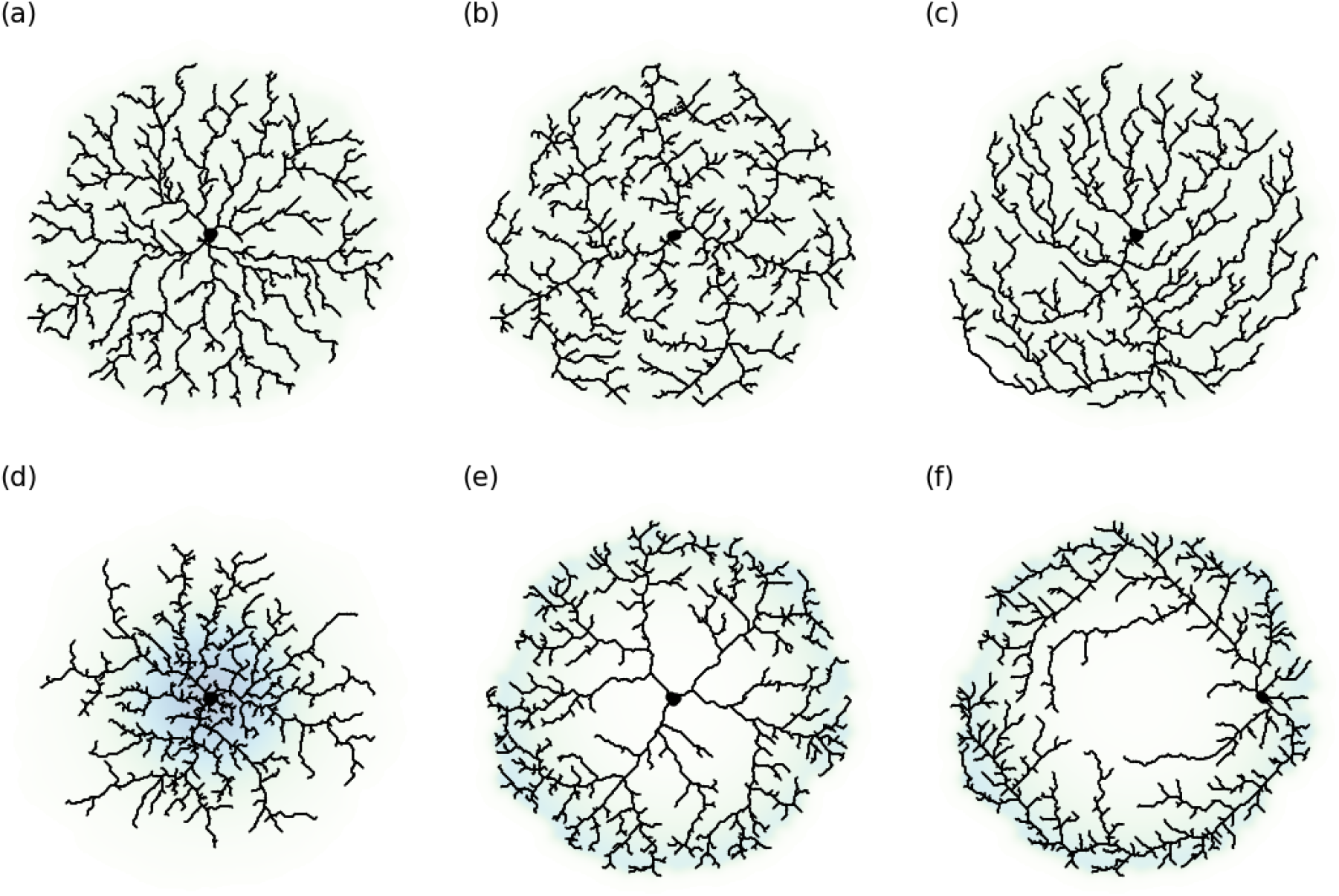
Effect of sampling strategies. Each tree is built by GD paths initiated at 500 seeds sampled with uniform (top row) and non-uniform (bottom row) probability density, shown as background color. GD paths are follow fast-marching travel times for a wave source marked with the black spot and located in the center in all cases, except for (f). Seed sampling order: (a) ordered by increasing distance from center, (b) ordered by decreasing distance from center, (c) ordered by decreasing distance from the central point at the top edge of the map, (d–f). In (f) the initial speed profile *v*_0_ was additionally weighted by the multiplication sampling probability 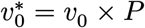 to make first paths to stay in the periphery resulting in a fundus-like morphology.

**Figure 4:**
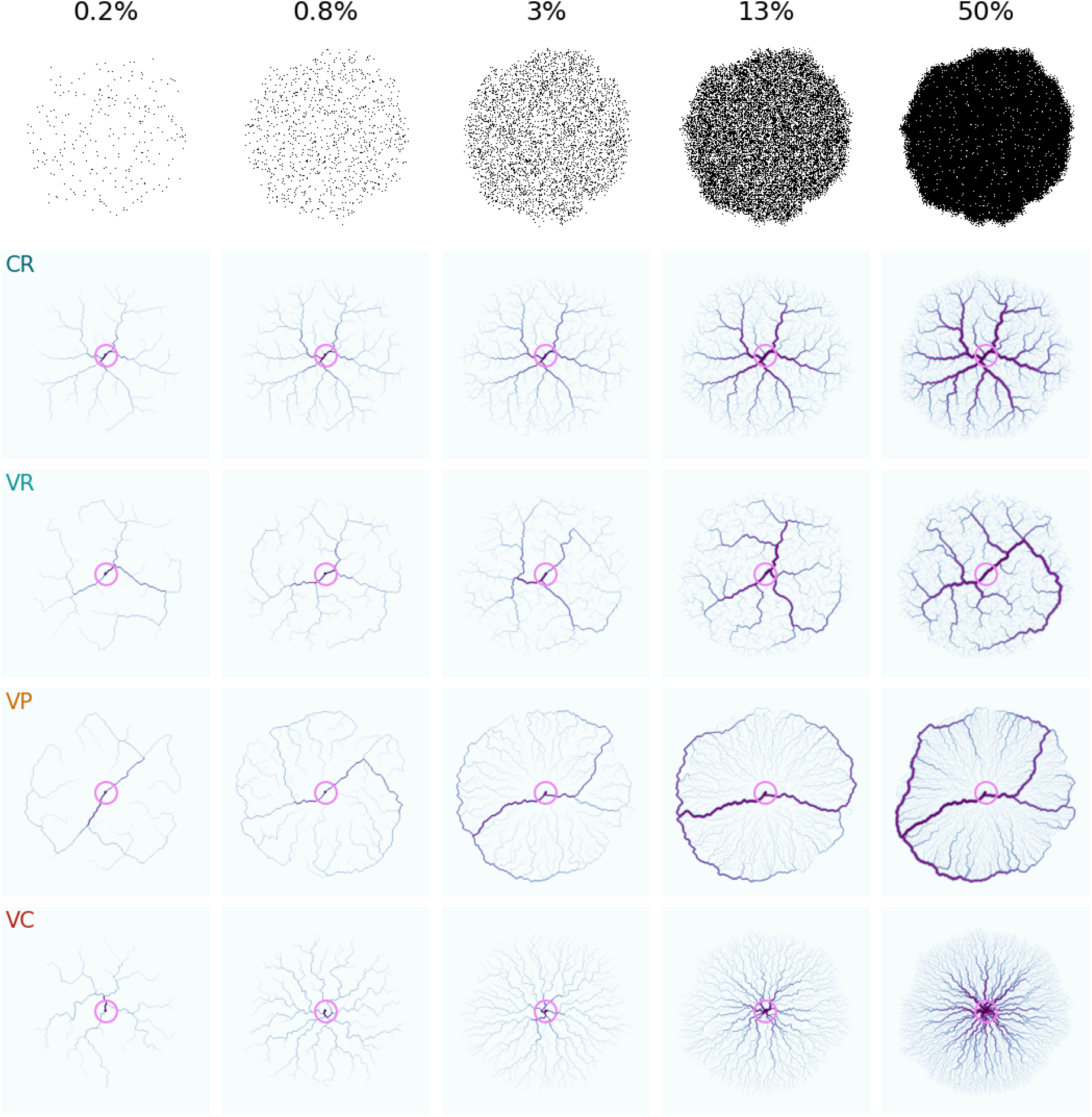
Effects of space filling and combination of seed ordering and targeting paths to young branches. Top row: illustration of seed densities (in percentage to total number of admissible pixels). CR, constant path target, random seed sampling order; VR, variable path target, random seed sampling order; VP, variable path target, sampling seeds from periphery to center; VC, variable path target, sampling seeds from center to periphery. Variable path targets are fast-marching wave sources originating from all nodes with occurrence count less than 32. Branch diameters in representations follow pipe flow model with *γ* = 2. Note less symmetric organization in VR, the “inside-out” morphology and VP and hypha-shaped morphology in VC.

## Results

### Effect of speed field on path merging and jitter

The proposed algorithm is based on tracing gradient descent (GD) paths in travel time maps, resulting form applying fast marching method to a field map with the wave source at tree root (or cell soma). Heterogeneity and smoothness of the speed field affect the resulting GD paths. If the speed profile is uniform, most of the paths are straight, rarely merge, and can run in parallel for considerable stretches (Figure 1 a). Adding high-pass Gaussian noise however makes paths more tortuous and less likely to run in parallel (Figure 1 b). Multi-scale colored noise leads to a more lobate trave time map and extends the spatial scale of path bends, while still allowing for some straight branches (Figure 1 c).

Spatial speed profile can be regarded as the relative likelihood of growing a branch from the root to a target point via one or another route. In the context of e.g. branched cell morphologies it seems not unnatural to consider local directionality of the process growth due to properties of extracellular matrix and space constraints from already existing cells. Applying a vesselness-enhancing filter, e.g. Sato, to a spatial field with independent Gaussian noise in each pixel leads to a characteristic sponge-like filamentous pattern, promoting anisotropic local correlations (Figure 1 d, top). When used as a speed field for fast-marching method, wave front follows these “fast-lane” filaments, leading to highly heterogeneous travel time map, while some locations are left occluded for the wave front access (Figure 1 d, middle). Gradient descent in such travel time map leads to highly jittered paths with increased probability of merging, due to wave front expansion “bottlenecks” (Figure 1 d, bottom). Multi-scale filamentous speed field, resulting from summation of Sato contrast images with different scale parameters applied to noisy input with independent Gaussian noise (Figure 1 e, top), induces a smoother travel time map with a small fraction of inaccessible regions (Figure 1 e, middle), and smoother GD paths (Figure 1 e, bottom). In this last variant the resulting branching pattern has qualitatively the most “natural” appearance and it is used as the starting speed field in the rest of the work.

### Effect of iterative speed field update by new branches

Another key component of the proposed algorithm is the path-speed feedback: newly built paths modify the speed field, reshaping travel-times and thus the subsequent path routes. Here we implement it as follows: each new path updates a count map of how many times each specific node was used to build a path; for converging paths this induces a hierarchy: nodes closer to a root will have been used more times and have higher count values than more distal ones. The resulting node-occurrence map *O* at iteration *i* can be added to the original speed profile after a rescaling transform *f* : *v*_*i*_ ← *v*_0_ + *f*(*O*_*i*_). When no update is made (*f*(·) → 0), the resulting tree, essentially, reflects a minimal wiring cost optimization, where wiring cost is defined by travel time profile (Figure 2, a). The choice of logarithmic *f*(*x*) → log *x*, linear *f*(*x*) → *kx* and power *f*(*x*) → *kx*^*α*^ scaling transforms makes already existing paths the “fast lanes” to reach the wave source, increasing the probability of path merging (Figure 2, b-d). In essence, this can be regarded as the conductance time optimization when connecting new distal seeds to a target root, resulting in trees at a balance between radially connected and minimum-spanning. Interestingly, the choice of update scaling only affects local branch attachment pattern details, but the general branching pattern remains similar in all three cases (Figure 2, b-d). Because the general pattern is not affected by the scaling transform, linear occurrence scaling will be used in the rest of the work.

### Effect of seed sampling strategy

So far we considered sparsely sampled seeds with random order and uniform probability. It is interesting to compare tree morphologies if the sampling probability density or order is non-uniform. To this end, we generated tree structures varying these parameters. In all simulations we considered trees build from 500 seed points, all paths converging to a single wave source by GD in travel time. Seed points were limited to the area with less than median travel time in the original speed profile, thus embedding morphologies to a disk with an irregular boundary and covering around half the modeling space area.

We first address the effect of seed sampling order and sample 500 random seed points with uniform probability within the allowed area (Figure 3 a-c). The tree branches are then sequentially built from these seed points in particular order. Starting with more central points and gradually increasing the distance between the seed and the wave source lead to a morphology similar to that obtained with random seed ordering (compare Figure 3 a to Figure 2 c). In contrast, first building paths from the most distant seeds and then gradually adding points closer to the root led to a distinct morphology pattern characterized by long distal processes going tangential to the admissible boundary and branch tips pointing inwards (Figure 3 b). Sorting seeds by decreasing distance to a point at the admissible boundary (top center in Figure 3 c) leads to an asymmetric morphology, displaying “branch taxis” towards the top part of the tree. This happens because the first few paths are built for the seed points approximately on the central axis and opposite to the reference location, and these paths go almost directly to the wave source at the center. Speed field update induced by the first paths makes travel times shorter along them, and subsequent GD paths from next sampled seeds are cued to converge on this central axis instead of directing straight to the wave source. This happens with a positive feedback until the new seeds start to appear in the top half of the modeling space, from where they are less likely to be affected by the travel time modification made by the early branches (Figure 3 c).

Next, we modify the density of sampled seeds while keeping the order of paths at random and compare morphologies with seeds preferentially sampled near the center and near the periphery of the modeling space (Figure 3 d–f). This also can have a mild ordering effect, as the seeds with higher probable locations are likely to crop up before the seeds with less probable locations, but the resulting morphologies were different from those obtained with ordered sampling (compare top and bottom rows of Figure 3). Under these conditions the denser seed sampling near the center leads to sparse processes in the periphery and denser coverage of the central region, reminiscent of gradual space colonization by a growing colony (Figure 3 e). On the other hand, higher seed density in the periphery leads to a spatial pattern similar to amacrine cells of the retina (Figure 3 e) with denser ramifications towards the outer boundary and less branching in the central region. Interestingly, shifting wave source (black point in the tree portraits) towards periphery and scaling initial speed field by the same periphery-enhancing spatial profile led to tree branching similar to fundus vasculature (Figure 3 f).

### Effect of dynamic path targets and space-filling properties

In the patterns above we considered morphologies with branches created from relatively sparse seeds, while the path target, i.e. wave source for the fast marching method, remained constant throughout the tree building process. Single path target naturally maps to e.g. a cell soma or tree root, but it also allows new paths to attach to nodes in existing branches irrespective of their occurrence count, i.e. the number of terminal points, whose paths go through the given node.

However in some contexts it would make sense to only allow new paths to attach to nodes in “young” branches, i.e. those with low path occurrence count. It seems also natural to treat the occurrence count as the number of terminal nodes supported by a branch node and link it to branch radius. Here we follow the pipe flow rule (Shinozaki et al., 1964), in which radius of a parent branch equals 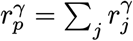, where *r*_*j*_ are its child branches. We use branch diameters for mainly for illustration purpose and optimize parameter *γ* to produce perceptually “natural” morphologies. Natural branching morphologies also vary in the density of the terminal branches, from relatively sparse in most neurons to practically space-filling in e.g. astrocytes. Below we observe how branching morphologies change with increasing density of seeding points, covering from 0.2% to 50% of all pixels in the models space as well as updated path target in combination with seed sampling ordering can produce dramatically different branching patterns (Figure 4).

We begin with branching morphologies built from increasingly dense seeds in the case of constant path target and random seeds sampling (Figure 4, CR). Increase in seed density leads to gradually more pronounced branch hierarchy, resulting in patterns resembling mammal astrocyte cells. Next we enforce path attachment only to “young” branches by using the nodes with small occurrence count as path targets, i.e. wave sources for fast marching method during the travel time update step. In this case, sampling path seed points at random leads to higher degree of competition between branches, resulting in a more asymmetric morphology with less primary branches stemming from the initial target location in the center (Figure 4, VR).

A combination of path target update and seed ordering further extends the morphology space accessible by the proposed algorithm. Presenting seeds ordered by decreasing distance to the center results in an “inside-out” morphology with a few thicker veins originating from the center, reaching the domain boundary and running along its circumference, while thinner processes reaching them from the inside in the centrifugal direction (Figure 4, VP). This happens because the first few paths must converge onto a central location, then the combination of speed profile update and path target update make the travel time along circumference smaller than on the inside of the domain. In contrast, presenting seeds ordered by increasing distance to the center leads to fanning-out structure of dense thin wrinkle-like processes with less pronounced diameter hierarchy than on other cases (Figure 4, VC). This later case can be mapped to a slow space colonization by e.g. a growing mold colony.

The morphologies resulting from the combinations of seed sorting and path target update differ in travel times from tip nodes to the center of the domain, i.e. initial path target (Figure 5, a). It has the most narrow distribution and the smallest mean for the structure with random seed sampling and constant path target at the center (CR), while targeting paths to young branches leads to wider travel time distributions with a larger mean (VR, VP, VC). Unexpectedly, the peak of the distribution is most shifted to the right for the morphology variant with path target update and center-first presented seeds, which is likely to be due to a lack of hierarchy and “fast-lane” branches with high wavefront speed in comparison to other structures. Travel times from the tip to the center can be also compared to geodesic path lengths, and the presented morphologies differ in how conduction scales with the wiring (Figure 5, a). The dependence is approximately linear for the constant target / random seed order (CR) and variable target/central ordering (VC) cases, with a noticeably steeper slope in the latter. In variable target / random seed order (VR) the scaling is a combination of linear part for short path lengths and nearly constant for longer path lengths, thus for a range of terminal nodes the travel time to the center doesn’t depend on the wiring length. Finally, the peculiar morphology of variable target / peripheral ordering (VP) induces a wide range travel times for most of the path lengths, reflecting the thin centrifugal processes running towards one of the circumferential major branches.

**Figure 5:**
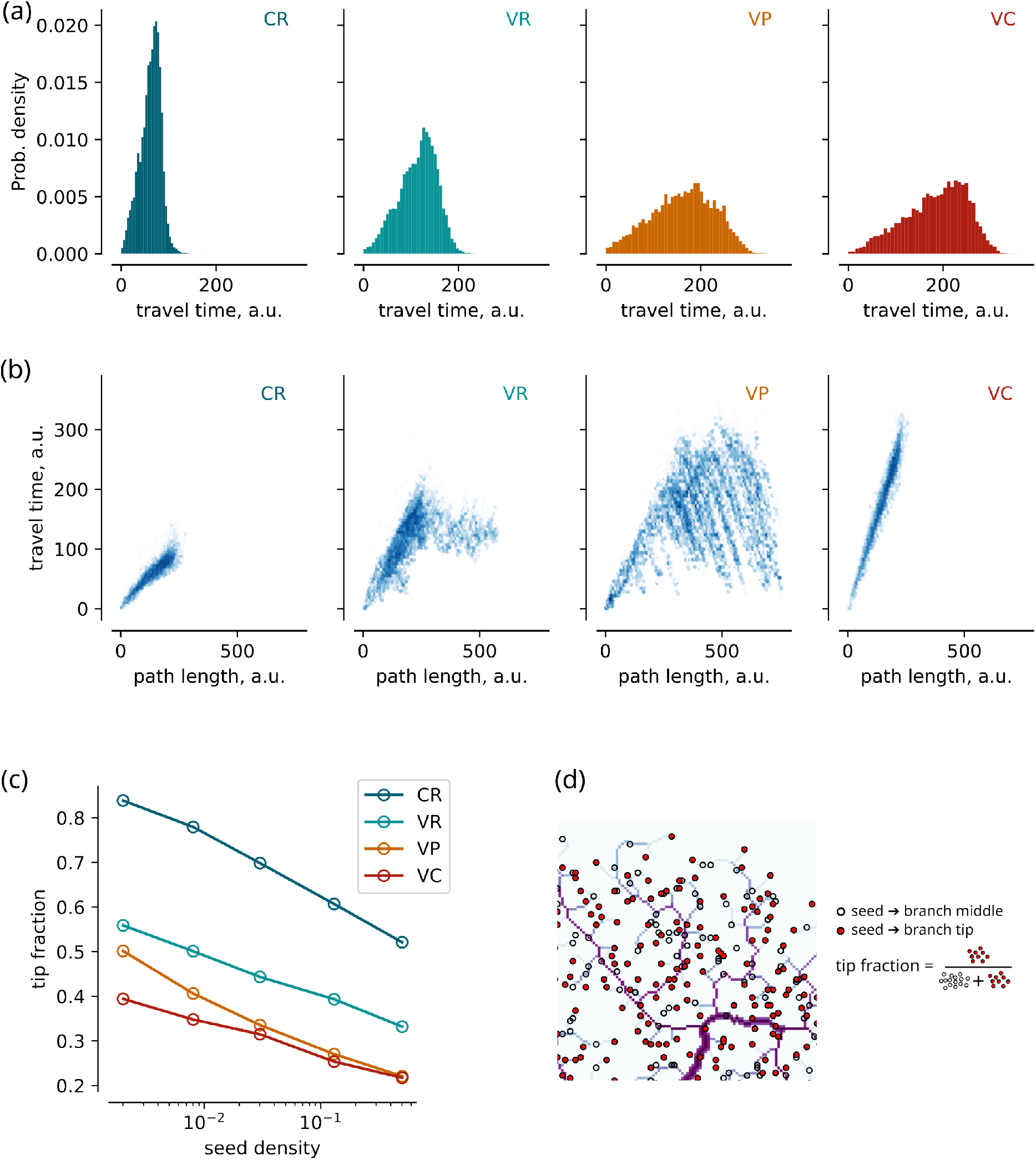
Effect of seed sampling and variable path target on efficiency of center-periphery communication. (a) Distributions of travel times from branch tips to the center of the domain for the morphology types presented in Figure 4 at 50% seed density; (b) scaling of of travel times with wiring path lengths for the same morphology types as in (a); note wider extent of path lengths in VR,VP and VC and less correlation between travel time and path length in VR and VP. (c) Tip fraction as a function of seed density for the same morphology types; note higher tip fraction in the CR. (d) Schematic representation of tip fraction calculation.

In the process of tree building some seed points will remain terminal nodes (“tips”), while some will become intermediate nodes if other new paths converge onto them. It is interesting to compare the fraction remaining “tips” to the total number of used seeding points between the morphology variants shown in Figure 4. This can be treated as a measure efficiency of the resulting tree — as in the number of targets subserved by a single source. Intuitively, using young branches as targets for path attachment should decrease the number of remaining terminal nodes, but whether this drop will be significant and the same for all seed ordering cases is not evident. To test this expectation, we calculated tip fractions for the four types morphologies shown in (Figure 4) at different seed densities (Figure 5, c). As expected, the case with constant target and random seed order (CR) displayed the highest tip-to-seed ratio, which gradually decreased with higher seed point densities. Preferential attachment to young branches (VR,VP,VC) reduced tip fraction, the least for random seed ordering and the most for center-first seed ordering. That is, the branching morphology in the latter case can support the smallest number of terminal nodes per tree size. On the other hand, more pronounced hierarchy in the other two cases allow for higher tip-to-seed ratio.

### Tiling networks and 3D extension

Simulation with a single path target leads to formation of a single connected tree. It is natural to extend the algorithm to multiple randomly placed wave sources for travel time calculation to serve as multiple competing targets for GD paths. This setting leads to a network of trees with a competitive space tiling, resembling astroglial networks in the hippocampus and neocortex (Figure 6,a) (Bushong et al., 2004). In this example random seed placement with 10% density and stationary path targets were used. Pipe flow model (Shinozaki et al., 1964) with *γ* = 1.85 was used to map node occurrence to local diameter, and color intensity corresponds to logarithm of node occurrence. Somata are placed artificially with a span corresponding to 0.5% percentile of time-travel map from each of the sources.

**Figure 6:**
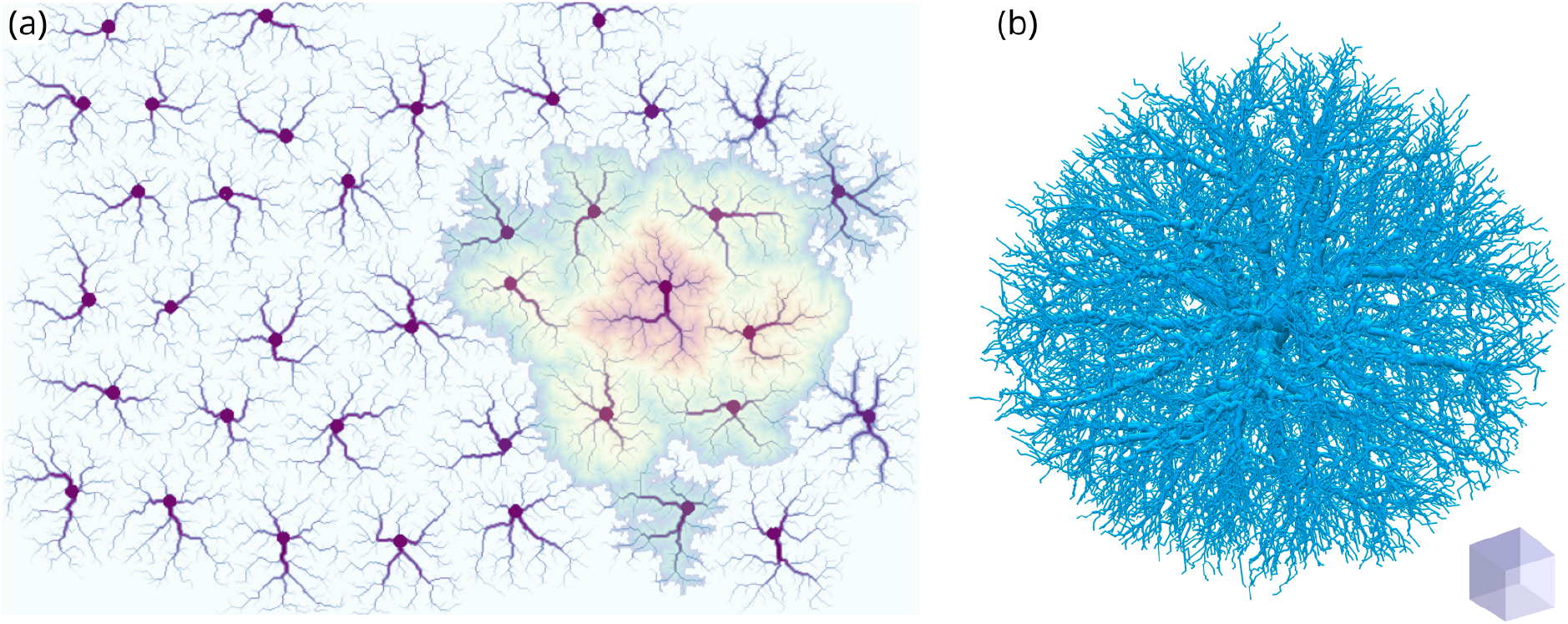
(a) a competitive tiling network of cells created with the proposed algorithm with 10% seeds density, branch diameters follow pipe flow model with *γ* = 1.85, colors from red to blue denote travel-time of a wave starting from a single cell soma and propagating with the speed proportional to the color intensity in the network image; (b) a three-dimensional tree, created with 10,000 randomly sampled seed points and single path target at the center. Diameters are based on pipe flow model with *γ* = 2.5

In the brain, neighboring astrocytes are connected by gap junctions (Anders et al., 2014; Houades et al., 2008), and ions and small molecules can redistribute across astrocytic network. Here we show the utility of the model in reproducing this network transport in the artificial astrocyte network by mapping travel time from a randomly chosen soma to the rest of the modeling domain with fast-marching wavefront speed proportional to pixel intensity (Figure 6, a). A wave is shown to propagate from domain to domain, first activating major branches and somata and next the rest of domain territories.

The proposed algorithm can also be directly extended to three dimensions, as demonstrated in (Figure 6), where GD paths from 10,000 randomly sampled seeding positions converged to a single central target. This opens a possibility to model three-dimensional branching structures over a wide range of morphology space.

## Discussion

We proposed a framework to simulate branching morphologies in “reverse time” as competitively merging paths following gradient descent (GD) in travel time maps. The latter are produced by wavefront expansion from a constant or updated source, which works like path target. The framework relies on 3 keypoints: (i) GD paths in travel time maps can attain a natural-looking jitter when the speed field is multi-scale filamentous, e.g. as produced by applying a vesselness filter with a range of spatial scales to random Gaussian noise images; (ii) the speed profile is updated by newly added paths to reshape travel time maps and subsequent gradient descent paths, increasing branch hierarchy in the resulting patterns; (iii) because the algorithm is essentially greedy, seed sampling order, especially in combination with dynamic path targets, allows to build an extensive range of morphologies.

Previous models of branching structures considered a problem of connecting point sources with a balance between constraints on wiring length and conductance delay (Chandrasekhar & Navlakha, 2019; Cuntz et al., 2010). Our algorithm can be interpreted in these terms as well. A gradient descent line in the travel time map is the shortest path between a seed point and a target (wave source), thus without speed profile update, the resulting tree will reflect the optimization for the wiring cost, allowing for a natural path tortuosity dictated by inhomogeneous speed field, while in direct tree-building algorithms jitter had to be added in post-processing (Cuntz et al., 2010). On the other hand, path-dependent speed profile update promotes preferential path convergence onto existing paths, increasing the weight of the conductance delay constraint. The latter can be further shaped by seed sampling probability of ordering and by setting the young branches as path targets during simulation.

The resulting morphologies can be tested for the efficiency of periphery to center communication and for the fraction of supported end terminals per seeding points used. Incidentally, the variant with a constant target at the center and random seed sampling turned out to be an optimal case in terms of supported terminals, while morphologies with preferential path attachment to young branches and random seeding demonstrated a flattened dependence of propagation time on wiring length. This links the current work to the optimal network flow models (Corson, 2010; Durand, 2007; Ronellenfitsch & Katifori, 2019), which also predict hierarchical tree-like structures to emerge in problems of resource distribution from a source to a set of sinks. A limitation of the proposed framework in comparison with the network flow model is that the latter can reproduce reticular morphologies in the case of resource fluctuations (Ronellenfitsch & Katifori, 2019), which gradient path merging model can’t do by construction. However, it seems interesting to explore a possibility of extending the proposed framework with an ability to create reticular morphologies.

Recent models of branching morphogenesis rely on a competing processes of tip elongation and tip branching (Hannezo et al., 2017; Ųcar et al., 2021), which basically links these models to a phenomenon of viscous fingering instability, in which the front of a less viscous liquid displacing a more viscous inmiscible liquid becomes unstable and fragments into branching structures, or it’s dual, diffusion-limited aggregation (Sarkar, 1985), (Bogoyavlenskiy, 2001). Essentially, merging of two gradient descent paths is equivalent to branching of a elongating tip, only simulated in a negative time direction, a “trick” used in dynamical systems theory and bifurcation analysis to explore unstable manifolds (Bosetti et al., 2010). Additionally, by varying the structure of the speed field, the proposed framework can be used to model branching morphologies intermediate between ballistic aggregation and diffusion-limited aggregation.

We hope that the algorithm regarded in this work can be useful to model realistic branching morphologies in computational biology, e.g. models of Ca^2+^ signaling or substance buffering in astrocytic networks. Some resulting morphologies look quite unusual, and may be of utility in design of biomimetic materials and biomorphs. Further development of the framework can include adapdations to create non-tiling networks with a variable extent of interdigitation, oriented speed profiles to mimic chemical cues in morphogenesis or the generalization of the structure-dependent path target updates to create self-moving patterns and model adaptive space exploration by hyphal networks or slime molds.

## Acknowledgements

This work was supported by the RSF grant # 23-44-00103

## Author contributions

A.K. worked on the algorithm implementation and prepared illustrations, T.L. worked on the algorithm implementation, A.B. conceived the idea and wrote the first draft, all authors worked on the manuscript text.

## Bibliography

Anders, S., Minge, D., Griemsmann, S., Herde, M. K., Steinhäuser, C., & Henneberger, C. (2014). Spatial Properties of Astrocyte Gap Junction Coupling in the Rat Hippocampus. Philosophical Transactions of the Royal Society of London. Series B, Biological Sciences, 369(1654), 20130600. 10.1098/rstb.2013.0600

Bogoyavlenskiy, V. A. (2001). Bridge from Diffusion-Limited Aggregation to the Saffman-Taylor Problem. Physical Review E, 63(4), 45305. 10.1103/PhysRevE.63.045305

Bosetti, H., Posch, H. A., Dellago, C., & Hoover, W. G. (2010). Time-Reversal Symmetry and Covariant Lyapunov Vectors for Simple Particle Models in and out of Thermal Equilibrium. Physical Review E, 82(4), 46218. 10.1103/PhysRevE.82.046218

Bushong, E. A., Martone, M. E., & Ellisman, M. H. (2004). Maturation of Astrocyte Morphology and the Establishment of Astrocyte Domains during Postnatal Hippocampal Development. International Journal of Developmental Neuroscience, 22(2), 73–86. 10.1016/j.ijdevneu.2003.12.008

Chandrasekhar, A., & Navlakha, S. (2019). Neural Arbors Are Pareto Optimal. Proceedings of the Royal Society B: Biological Sciences, 286(1902), 20182727. 10.1098/rspb.2018.2727

Corson, F. (2010). Fluctuations and Redundancy in Optimal Transport Networks. Physical Review Letters, 104(4), 48703. 10.1103/PhysRevLett.104.048703

Cuntz, H., Forstner, F., Borst, A., & Häusser, M. (2010). One Rule to Grow Them All: A General Theory of Neuronal Branching and Its Practical Application. Plos Computational Biology, 6(8), e1000877. 10.1371/journal.pcbi.1000877

Durand, M. (2007). Structure of Optimal Transport Networks Subject to a Global Constraint. Physical Review Letters, 98(8), 88701. 10.1103/PhysRevLett.98.088701

Garrido, S., Alvarez, D., & Moreno, L. E. (2020). Marine Applications of the Fast Marching Method. Frontiers in Robotics and AI, 7, 2. 10.3389/frobt.2020.00002

Garrido, S., Moreno, L., Martín, F., & Álvarez, D. (2017). Fast Marching Subjected to a Vector Field–Path Planning Method for Mars Rovers. Expert Systems with Applications, 78, 334–346. 10.1016/j.eswa.2017.02.019

Hannezo, E., Scheele, C. L., Moad, M., Drogo, N., Heer, R., Sampogna, R. V., Van Rheenen, J., & Simons, B. D. (2017). A Unifying Theory of Branching Morphogenesis. Cell, 171(1), 242–255. 10.1016/j.cell.2017.08.026

Houades, V., Koulakoff, A., Ezan, P., Seif, I., & Giaume, C. (2008). Gap Junction-Mediated Astrocytic Networks in the Mouse Barrel Cortex. The Journal of Neuroscience, 28(20), 5207–5217. 10.1523/JNEUROSCI.5100-07.2008

Parekh, R., & Ascoli, G. (2013). Neuronal Morphology Goes Digital: A Research Hub for Cellular and System Neuroscience. Neuron, 77(6), 1017–1038. 10.1016/j.neuron.2013.03.008

Ronellenfitsch, H., & Katifori, E. (2019). Phenotypes of Vascular Flow Networks. Physical Review Letters, 123(24), 248101. 10.1103/PhysRevLett.123.248101

Runions, A., Lane, B., & Prusinkiewicz, P. Modeling Trees with a Space Colonization Algorithm. 9.

Salmon, C. K., Syed, T. A., Kacerovsky, J. B., Alivodej, N., Schober, A. L., Sloan, T. F., Pratte, M. T., Rosen, M. P., Green, M., Chirgwin-Dasgupta, A., Mehta, S., Jilani, A., Wang, Y., Vali, H., Mandato, C. A., Siddiqi, K., & Murai, K. K. (2023). Organizing Principles of Astrocytic Nanoarchitecture in the Mouse Cerebral Cortex. Current Biology, 33(5), 957–972. 10.1016/j.cub.2023.01.043

Sarkar, S. K. (1985). Saffman-Taylor Instability and Pattern Formation in Diffusion-Limited Aggregation. Physical Review a, 32(5), 3114–3116. 10.1103/PhysRevA.32.3114

Sato, Y., Nakajima, S., Shiraga, N., Atsumi, H., Yoshida, S., Koller, T., Gerig, G., & Kikinis, R. (1998). Three-Dimensional Multi-Scale Line Filter for Segmentation and Visualization of Curvilinear Structures in Medical Images. Medical Image Analysis, 2(2), 143–168. 10.1016/S1361-8415(98)80009-1

Sethian, J. A. (1996). A Fast Marching Level Set Method for Monotonically Advancing Fronts. Proceedings of the National Academy of Sciences, 93(4), 1591–1595. 10.1073/pnas.93.4.1591

Shinozaki, K., Yoda, K., Hozumi, K., & Kira, T. (1964,). A Quantitative Analysis Of Plant Form-The Pipe Model Theory : I.Basic Analyses (Issue 3). The Ecological Society of Japan. 10.18960/seitai.14.3_97

Ųcar, M. C., Kamenev, D., Sunadome, K., Fachet, D., Lallemend, F., Adameyko, I., Hadjab, S., & Hannezo, E. (2021). Theory of Branching Morphogenesis by Local Interactions and Global Guidance. Nature Communications, 12(1), 6830. 10.1038/s41467-021-27135-5

